# TRPV1 regulates opioid analgesia during inflammation

**DOI:** 10.1101/274233

**Authors:** Lilian Basso, Reem Aboushousha, Churmy Yong Fan, Mircea Iftinca, Helvira Melo, Robyn Flynn, Francina Agosti, Morley D. Hollenberg, Roger Thompson, Emmanuel Bourinet, Tuan Trang, Christophe Altier

## Abstract

Acute inflammation in humans or mice enhances the analgesic properties of opioids. However, the inflammatory transducers that prime opioid receptor signaling in nociceptors are unknown. We found that TRPV1^−/−^ mice are insensitive to peripheral opioid analgesia in an inflammatory pain model. We report that TRPV1 channel activation drives a MAPK signaling pathway accompanied by the shuttling of β-arrestin2 to the nucleus. This shuttling in turn prevents: β-arrestin2-receptor recruitment, subsequent internalization of agonist-bound mu opioid receptor (MOR), and suppression of DAMGO-induced inhibition of N-type calcium current observed upon desensitization. Consequently, inflammation-induced activation of TRPV1 preserves opioid analgesic potency in a mouse model of opioid receptor desensitization. Overall, our work reveals a TRPV1-mediated signaling mechanism, involving β-arrestin2 nuclear translocation, that underlies the peripheral opioid control of inflammatory pain. Our data single out TRPV1 channels as modulators of opioid analgesia.

---

Pain modulation and inflammation are two integrated biological responses that function in cooperation following injury. Particularly known for their analgesic properties, opioids target receptors expressed throughout the afferent pain pathway, including central and peripheral nerve terminals of primary afferent neurons^1-3^. Indeed, local endogenously produced opioids, β-endorphin, enkephalins, and dynorphins exert efficient inhibitory control of pain at both spinal^1,2^ and peripheral sites following inflammation^4-8^. At inflammatory sites, immune-derived opioids are released to counteract pain effectively and facilitate healing. Moreover, alteration in the endogenous opioid system in the spinal cord was recently proposed to prevent the transition from acute inflammatory to chronic pain^1^, particularly through tonic μ-opioid receptor (MOR) activity that suppresses hyperalgesia following resolution of inflammation. Thus, both endogenous and xenobiotic opioids are most effective immediately after tissue injury, suggesting that the early phases of inflammation primes opioid receptor signaling in primary afferent neurons^7,9^. Alterations in this priming might contribute to a decrease in the therapeutic window of opioids, requiring increased analgesic doses that can lead to side effects.

At sites of acute local inflammation in humans or mice, the analgesic effect of opioids is enhanced^10^, whereas in contrast, the application of opioids to a human peripheral nerve plexus does not elicit analgesia in non-inflamed tissue^11^. At these sites, TRPV1 is the main endpoint target of inflammatory mediators, including reactive-oxygen species, metabolites of polyunsaturated fatty acids, inflammatory acidosis or toxins released by pathogen^12^. Moreover, both the activity and trafficking of the channel are sensitized by GPCR-targeted mediators (BK, PGE2, proteases)^13-16^, which in turn causes hyperalgesia during tissue inflammation^17^ and in many immune-associated diseases^12,18-21^.

Along these lines, recent findings suggest that MORs expressed in TRPV1^+^ nociceptors are essential contributors to analgesic tolerance^22^ yet, the molecular mechanisms underlying these biological regulations remain unknown. We set out to test whether the TRPV1 channel, a central integrator of inflammatory signals^12^, could prime opioid receptor signaling or modulate opioid analgesia during inflammation. Here, using the complete Freund’s adjuvant (CFA) inflammatory pain model, we show that TRPV1^−/−^ mice exhibit alterations in both endogenous and exogenous opioid analgesia. We reveal further that, upon TRPV1 activation, β-arrestin2, a critical regulator of opioid signaling, is routed to the nucleus through a calcium-dependent mitogen-activated protein kinase (MAPK) signaling pathway. As a result of this exclusion of β-arrestin2 from the cytosol, activated MORs are unable to recruit β-arrestin2 and internalize during agonist-induced desensitization. This reduced MOR desensitization has major implications for opioid-induced analgesia and β-arrestin2 biased signaling *in vivo*. Overall, our data uncover a mechanism by which inflammation-mediated activation of TRPV1 channels minimizes opioid receptor opioid receptor desensitization to optimize analgesia. This TRPV1-opioid receptor interaction could be pharmacologically targeted in clinical settings to improve pain management by opioids.

## Results

### TRPV1 regulates endogenous opioid-mediated analgesia during inflammation

To determine the molecular link between inflammation and opioid analgesia we asked if, as a central integrator of inflammatory signals, TRPV1 channel could modulate intrinsic opioid receptor function during peripheral inflammation. Therefore, we assessed inflammation-induced endogenous opioid antinociception in TRPV1^−/−^ mice. TRPV1 is likely the best characterized sensory transducer that contributes to thermal hyperalgesia in acute models of inflammation^23-25^, although the channel is expressed in polymodal C fibers that process both thermal and mechanical hyperalgesia during long-lasting inflammation^26^. Using the CFA model of chronic inflammatory pain, we evaluated peripheral immune-derived endogenous opioid analgesia^6,27-30^ by measuring paw withdrawal threshold (PWT) during the later phase of inflammation. In WT animals, PWT was decreased by 50% at day 4 after intraplantar injection of CFA, and animals started to recover at day 6. Treatment with naloxone-methiodide (Nal-M), a peripherally restricted non-selective and competitive opioid receptor antagonist, delayed the recovery time as attested by the increased PWT from day 6 to day 10 in both male (**Fig. 1a and b**) and female mice (**Supplementary Fig. 1a and b)**. In TRPV1^−/−^ mice, we found that PWT was decreased by 40% at day 4, and then recovered gradually until day 14. Strikingly, treatment with Nal-M in both TRPV1^−/−^ male (**Fig. 1a and b**) and female mice (**Supplementary Fig. 1a and b**), had no effect on PWT at any time of the treatment. This absence of endogenous opioid analgesia in TRPV1-/- mice was due neither to a difference in inflammation (edema was similar between WT and KO mice), nor the capacity of pain-modulating immune cells to produce endogenous opioids and infiltrate the inflamed paw at day 8 of CFA (**Supplementary Figure 2a-c**). Thus, as previously reported, local opioid signaling is fine-tuned by inflammation^29^, and our results indicate that TRPV1 is a central contributor to this process.

**Figure 1.**
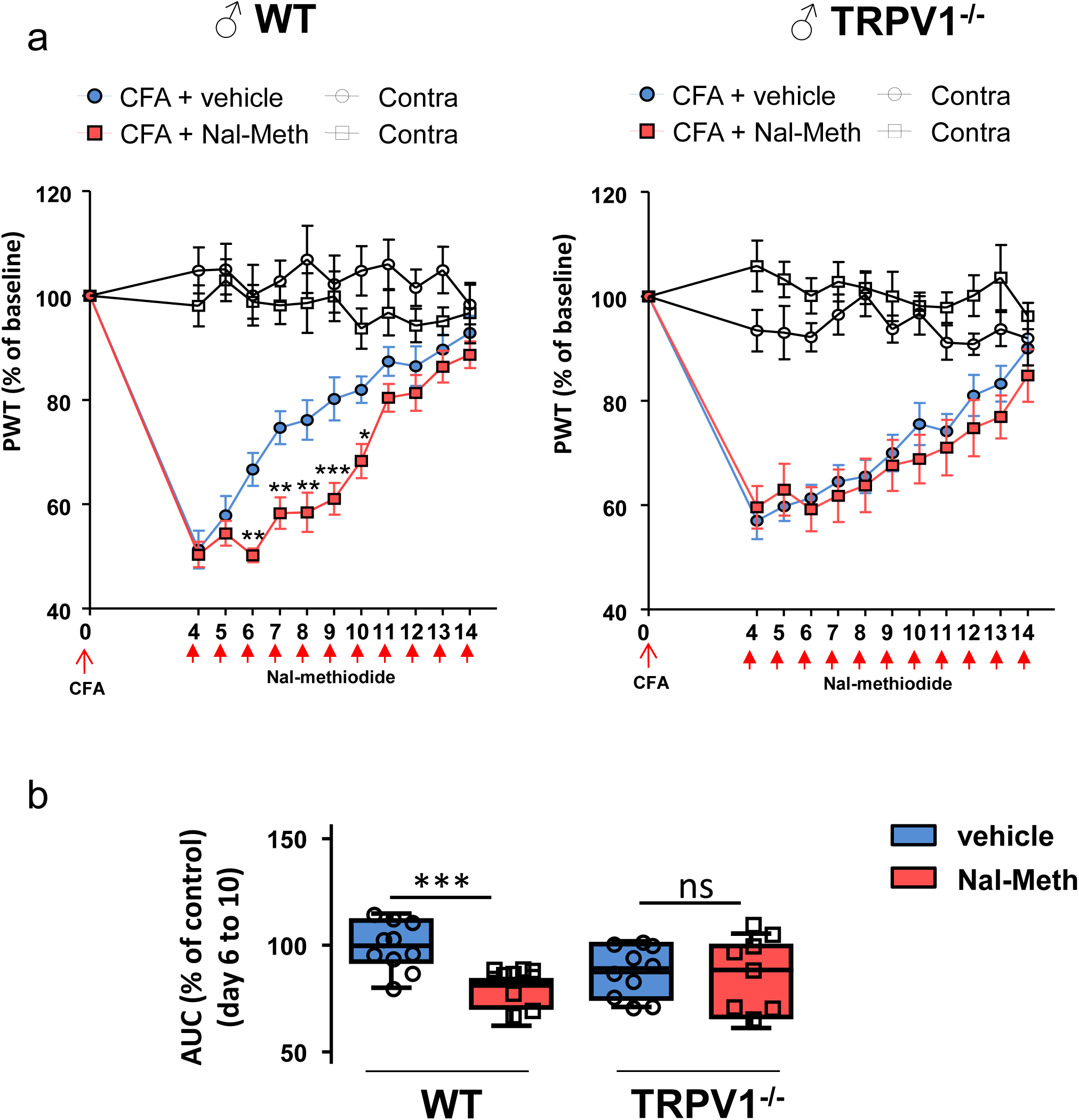
Peripheral endogenous control of inflammatory pain is absent in TRPV1 deficient mice. CFA (30μl; 1mg/ml) was injected into the plantar surface of the right hind paw of WT (left graph) or TRPV1 -/- mice (right graph). Mechanical hypersensitivity was assessed by measuring paw withdrawal thresholds (PWT) using a dynamic plantar aesthesiometer. Mice were injected with either 10µl PBS (blue circle; n=10 for each genotype) or Naloxone-Methiodide (2mg/ml, red square; n=10 for WT and n=8 for TRPV1-/-) in the ankle from day 4 to day 14, 30 minutes before mechanical nociception assessment. (a) Data are expressed as the mean ±SEM of PWT measured in the inflamed (CFA) ipsilateral paw (color) and contralateral saline-injected paw (grey) normalized to baseline. (* p < 0.05; ** p < 0.01; *** p < 0.001; two-way ANOVA followed by Bonferroni post hoc test). Area under the curve between day 6 and 10 were calculated and expressed as % of vehicle treated animals for each genotype. Data are expressed as the mean min-max (*** p < 0.001; Mann-Whitney U test) (b)

### TRPV1 activation disrupts MOR-β-arrestin2 interaction

To delineate how opioid modulation is impaired in TRPV1^−/−^ mice, we focused on β-arrestin2, a central hub signaling protein that modulates both opioid receptor desensitization and recycling^31^. A crosstalk between β-arrestin2 and the TRPV1 channel has recently emerged, particularly in the context of opioid receptor sequestration^32,33^. To determine whether activation of TRPV1 redirects β-arrestin2 away from the agonist-bound MOR, we tested the effect of capsaicin using a BRET assay that monitored interaction between MOR-Rluc8 and β-arrestin2-Venus. As shown in **Fig. 2**, DAMGO-mediated activation of MOR induces the recruitment of β-arrestin2 to MOR, indicated by the increase in BRET signals over time. In the absence of TRPV1, treatment with capsaicin does not induce a BRET response or affect the DAMGO-induced recruitment of β-arrestin-2 to MOR. However, in TRPV1 expressing cells, capsaicin co-treatment along with DAMGO completely blocks the interaction between MOR and β-arrestin2 (**Fig. 2b and c**), at low and even saturating concentrations of DAMGO (**Fig. 2d**), while capsaicin alone had no effect. Use of the reversed pair of BRET biosensors (βARR2-Rluc + MOR-YFP) yielded to similar results (**Supplementary Fig. 3**), thus confirming the inhibitory effect of TRPV1 activation on MOR/β-arrestin2 interaction. Interestingly, As shown in **Fig. 2**, co-immunoprecipitation of MOR with TRPV1 indicates the formation of a receptor-channel signaling complex at steady state (**Fig. 2e**), and the saturating BRET signals obtained for MOR and TRPV1 co-expression suggests a direct interaction between MOR and TRPV1 in HEK cells, with or without DAMGO (**Fig. 2f**). Therefore, our findings suggest that proximity of the TRPV1 channel to MOR, may alter the β-arrestin2-biased agonism of MOR by redirecting β-arrestin2 or preventing its recruitment to the receptor. Interestingly, we also found that this mechanism applies to other neuronal GPCRs, as the Proteinase-activated receptor-2 (PAR2), which like the MOR is involved in inflammatory pain, showed impaired β-arrestin2 recruitment following capsaicin co-treatment. Given that β-arrestin2 is an essential modulator of MOR desensitization and recycling, we next tested the functional outcome of TRPV1 activation on receptor internalization. Using confocal microscopy we found that, as extensively reported, exposure to DAMGO for 20 min promoted significant MOR internalization in HEK cells transfected with MOR-YFP and TRPV1-mCherry. In contrast, cells co-stimulated with DAMGO and capsaicin (or RTX, not shown), had significantly less internalized receptor **(Fig 3a and b**). Importantly, deletion of β-arrestin2 using a CRISPR β-arrestin2 knockout cell line **(Supplementary Fig. 4**), eliminated DAMGO-evoked internalization, and thus mimicked the DAMGO+capsaicin condition in β-arrestin2 WT cells. **(Fig. 3 c and d**).

**Figure 2.**
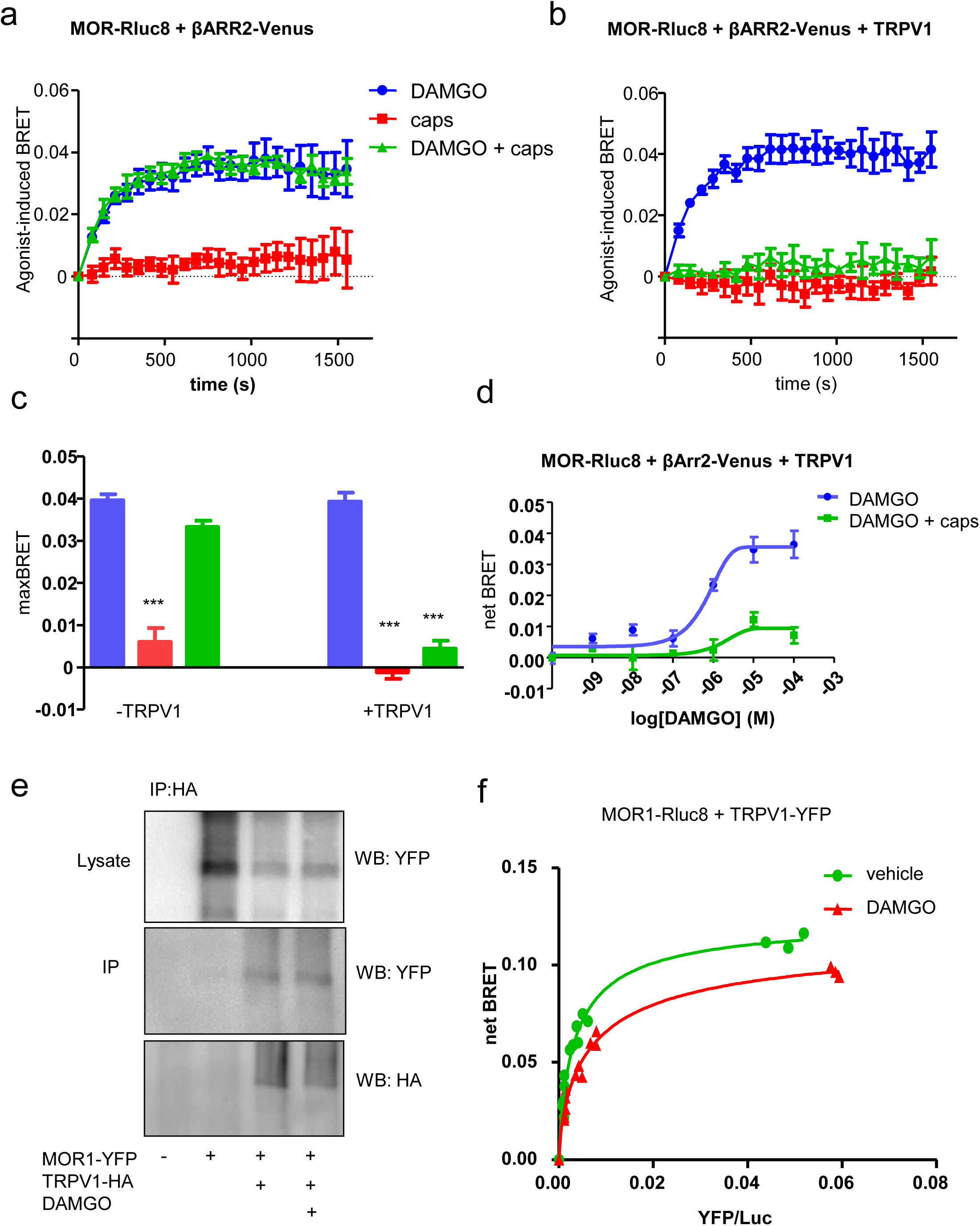
Activation of TRPV1 prevents MOR - β-arrestin2 interaction upon agonist stimulation. Time course of agonist-induced BRET (DAMGO=10μM, capsaicin=500nM) between MOR-Rluc8 and β-ARR2-Venus in absence a) or presence b) of TRPV1 channel. c) Quantification of the maximum BRET signal obtained in a) and b). Data represent the mean± S.E.M of the last four time points on the time course curve. d) Dose-response curve of the net BRET between MOR-Rluc8 and β-arrestin2-Venus in TRPV1-transfected cells following DAMGO or DAMGO+capsaicin exposure. (***P<0.01, N=4 experiments). e) Co-immunoprecipitation of TRPV1-HA with MOR-YFP in transfected HEK cells. f) Saturation BRET curve between MOR1-Rluc8 and TRPV1-YFP in transfected HEK cells treated or not with DAMGO.

**Figure 3.**
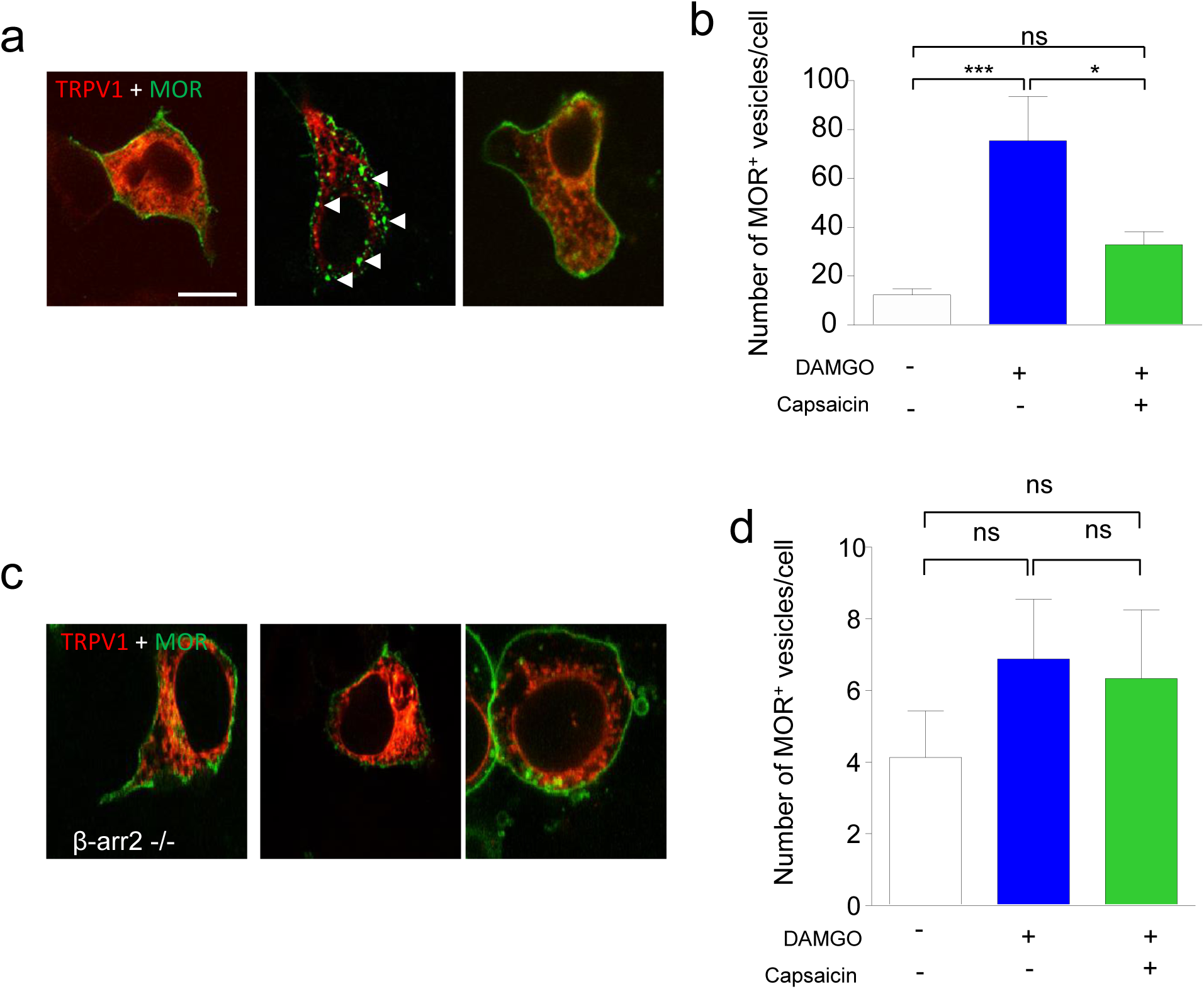
Activation of TRPV1 prevents MOR internalization upon agonist stimulation. WT (a) or β-arrestin2 KO (c) HEK cells were transfected with MOR-GFP, TRPV1-mCherry and untreated (WT HEK n=16 / β-arr-2 KO HEK n=8), or treated with either DAMGO alone (1µM; WT HEK n=14 / β-arr-2 KO HEK n=16) or DAMGO and Capsaicin (1µM; WT HEK n=29 / β-arr-2 KO HEK n=12). Number of vesicles per cell was counted using ImageJ software and presented in (b) and (d). Results from two independent experiments are expressed as mean±SEM (* p < 0.05;*** p < 0.001, Kruskal-Wallis test followed by Dunn’s post hoc test when appropriate).

### TRPV1 activation induces both β-arrestin2 nuclear localization and ERK activation

As Por et al. reported that β-arrestin2 could interact with TRPV1 upon channel stimulation^32^, we asked whether channel activation could, through competitive interaction, prevent the recruitment of β-arrestin2 to MOR. Using our BRET assay, we found no evidence for an interaction between βarr2-YFP and TRPV1-Rluc (**Fig. 4a and b**) either at steady state, or upon channel activation by the potent TRPV1 ligand RTX. In contrast, BRET signals indicate a strong interaction between the Rluc-TRPV1 and YFP-TRPV1, confirming the functional assembly of tagged-subunits into oligomeric channels. As β-arrestin-2 does not interact with either MOR or TRPV1 upon capsaicin exposure, we examined the fate of β-arrestin2-YFP using spinning disk confocal microscopy on TRPV1 transfected cells. We made the striking observation that channel activation promotes a rapid translocation of β-arrestin2 to the nucleus (**Fig. 4c**). Quantification of β-arrestin2 nuclear localization indicates that >70% of TRPV1mCherry-expressing cells display nuclear translocation of β-arrestin2 at 15 min. after RTX application (**Fig. 4d and e**). This effect was slowly reversed upon RTX wash out and did not occur in absence of the channel (**Supplementary Fig. 5a and b**). Nuclear Hoechst blue staining confirmed the sequestration of β-arrestin2 into the nucleus after RTX challenge (**Fig. 4f**), an effect that could also be observed with capsaicin, using confocal imaging (**Fig. 4g**), or assessed by bystander BRET with the renilla GFP fused to the nuclear localization sequence (NLS) PKKKRKVEDPKS targeting the nucleus (**Fig. 4h**)^34^. In contrast, activation of TRPA1, another calcium-permeant member of the TRP channel family, did not trigger β-arrestin2 translocation **(Supplementary Fig. 5c**). As for the activation of TRPA1, increasing intracellular calcium concentrations using the ionophore, ionomycin, also failed to stimulate the nuclear translocation of β-arrestin2 (not shown). Importantly, immunostaining of β-arrestin2 in TRPV1-expressing neurons identified using a TRPV1-YFP reporter mouse indicated similar results (**Fig. 4i**). Thus, activation of TRPV1 rapidly relocates β-arrestin2 into the nucleus both in expression systems and in native sensory neurons. As a consequence, β-arrestin2 that normally colocalizes with internalizing MOR-containing vesicles, is now sequestered within the nucleus upon channel stimulation. This sequestration prevents agonist-induced internalization of the MOR **(Supplementary Fig. 6**).

**Figure 4.**
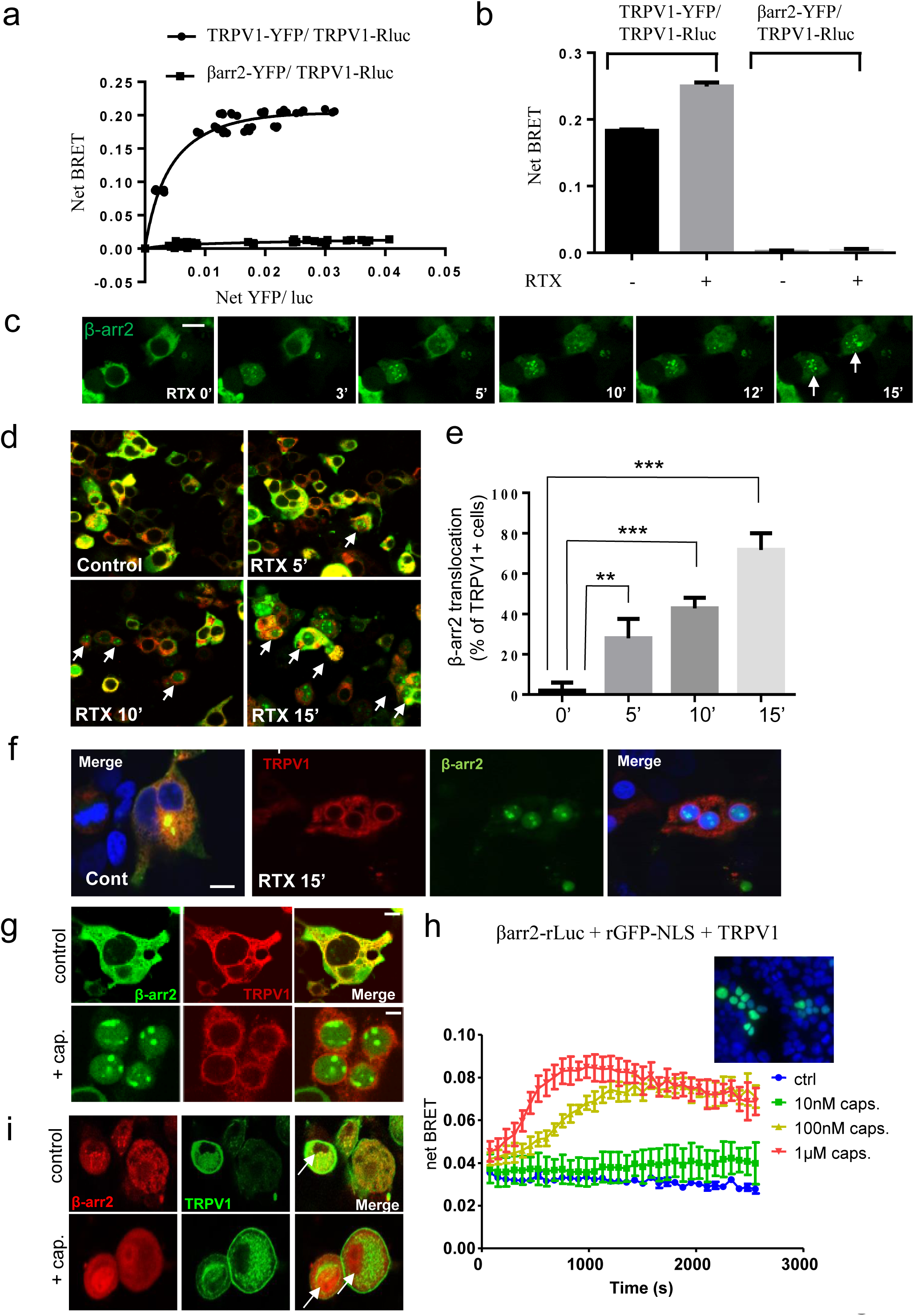
Activation of TRPV1 mediates translocation of β-arrestin-2 to the nucleus. (a) Saturation BRET curve between β-arrestin2-YFP and TRPV1-Rluc or TRPV1-YFP and TRPV1-Rluc as a positive control. (b) NetBRET between TRPV1-YFP and TRPV1-Rluc or β-arrestin-2-YFP and TRPV1-Rluc upon RTX treatment. (c) Spinning disk confocal images of HEK cells transfected with TRPV1 channel and β-arrestin2-YFP in response to bath applied RTX (10 nM) at 37°C for different time points (Scale bar = 10 µm). (d) Representative confocal images showing β-arrestin-2-GFP (green) nuclear translocation in HEK transfected with TRPV1-mCherry (red) after exposure to RTX (10 nM). (Scale bar = 20 µm). (e) Percentage of TRPV1-mCherry positive HEK cells showing β-arrestin2 nuclear translocation following TRPV1 activation (10 nM) at 5, 10 or 15 min of RTX treatment (n = 10 fields per condition from 3 separated transfections, RTX 0 min 2% ± 2, RTX 5 min 27.9% ± 7.4, RTX 10 min 42.7% ± 6.9, RTX 15 min 71.7% ± 7.24. Data are expressed as mean values ± SEM. p: ** <0.001, *** <0.0001; unpaired t-test.) (f) Nuclear Hoechst blue staining confirming β-arrestin-2 (green) translocation to the nucleus in response to activation of TRPV1-mcherry (red) by RTX (10 nM, 15 min). (Scale bar = 10 µm). (g) Confocal images of cells transfected with TRPV1-mCherry (red) and β-arrestin2-YFP (green). Treatment with capsaicin (1 µM, 15 min) also induced β-arrestin-2 translocation into the nucleus. (Scale bar = 10 µm). (h) Bystander BRET measured over time in HEK cells expressing TRPV1, β2ARR-RlucII and renilla GFP-NLS. Capsaicin was injected a t=0. Inset image shows the nuclear localization of rGFP-NLS (500ng transfected) in cells co-stained with DAPI. (i) Immunolabelling of β-arrestin-2 (red) in TRPV1-expressing DRG neurons (green) isolated from Ai32/TRPV1-cre mouse. Neurons were stimulated with capsaicin (100 nM) for 10 min. Note the nuclear translocation of β-arrestin2 in TRPV1(+) neurons.

As arrestins are important scaffolds for the mitogen-activated protein kinase (MAPK) signaling pathway, we tested the effect of TRPV1 stimulation on extracellular signal-regulated kinases (ERK1/2). In TRPV1 transfected cells, RTX induces a transient phosphorylation of ERK1/2 at 5 min. post-stimulation (**Fig. 5 a and b**). ERK1/2 activation is blocked by chelating extracellular calcium with EGTA (10 mM) (**Fig. 5c**), as is the β-arrestin2 nuclear translocation (**Fig. 5d**), confirming that calcium influx through activated TRPV1 channels mediates both ERK1/2 activation and β-arrestin2 translocation. Several kinases have been implicated upstream of ERK1/2 activation, including Src and different isoforms of PKC^35,36^. We found that pharmacological inhibition of PKCβII by CGP53353 blocks ERK1/2 phosphorylation, whereas selective inhibition of the α or βI isoforms with the GF109203X compound did not (**Fig. 5 e and f**). Accordingly, TRPV1 activation by RTX directly activates PKCβII (**Fig. 5g**). Finally, to confirm that β-arrestin2 was a central chaperone in the ERK signaling cascade engaged by TRPV1 stimulation, we tested the effect of knocking out β-arrestin2 in capsaicin-stimulated TRPV1 cells. Western blot analysis indicated an absence of ERK1/2 activation at 5 min. of TRPV1 stimulation on cells treated with β-arrestin2 but not scrambled siRNA (**Fig. 5h**). Altogether, these findings indicate that activation of the TRPV1 channel engages a signaling cascade involving both a Ca2^+^ and PKCβII-dependent ERK1/2 activation, which coincides with the translocation of β-arrestin2 into the nucleus.

**Figure 5.**
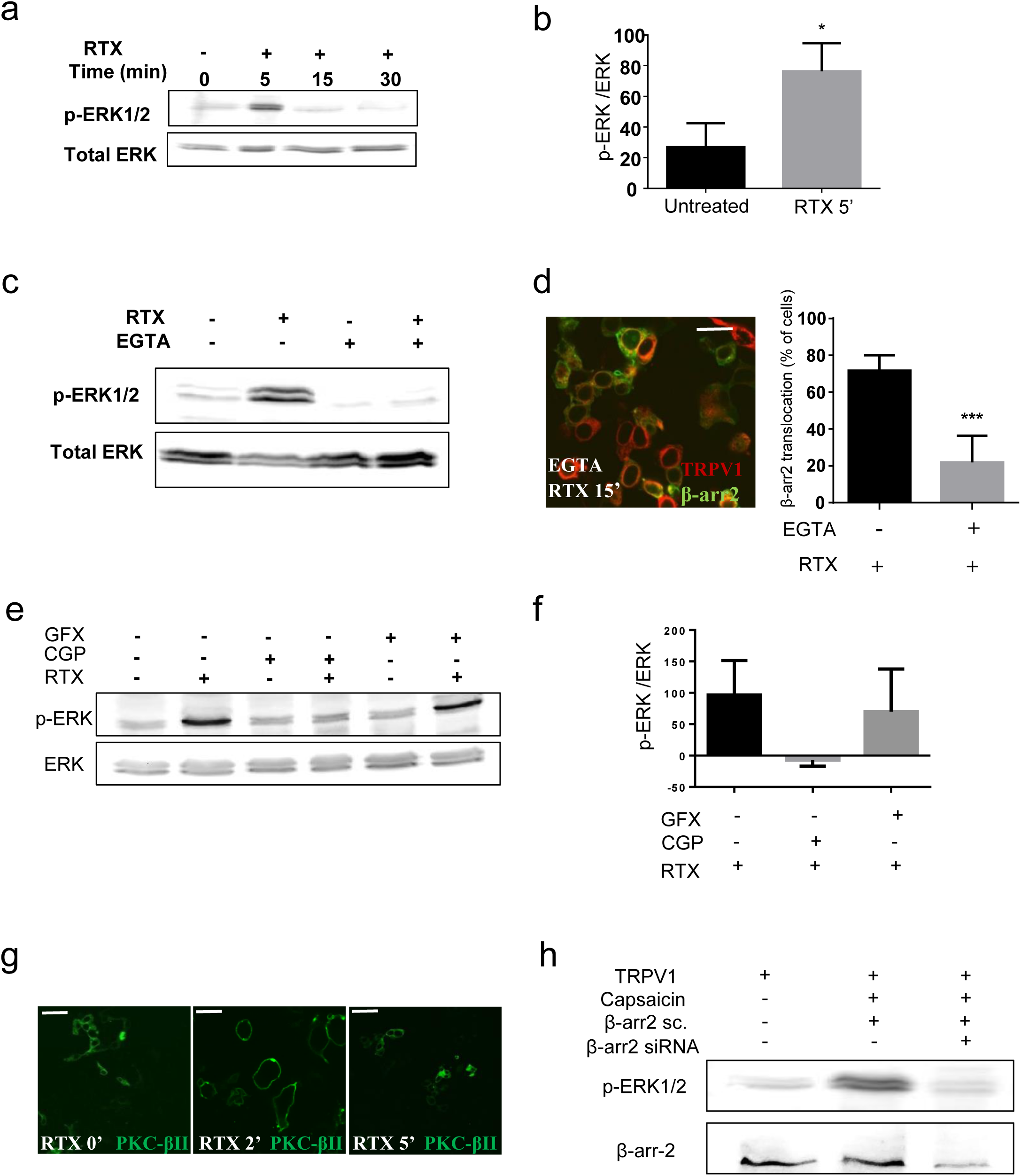
Activation of TRPV1 induces transient ERK1/2 phosphorylation. (a) Representative Western Blot of pERK1/2 and total ERK in response to TRPV1 activation by RTX (10 nM) in transfected HEK cells. (b) Densitometry analysis of pERK1/2 normalized to total ERK at 5 min post RTX treatment (Unt: 26.1 ± 14, RTX: 79 ± 13; P*<0.05; n = 3). (c) Effect of the Ca2+ chelator EGTA (10 mM) on RTX-induced pERK1/2 probed by western blot. (d, left) Confocal image of TRPV1-mCherry/β-arrestin-2-YFP transfected cells exposed to RTX (10 nM, 15 min) in the presence of EGTA (10 mM). Note that β-arrestin2 did not display nuclear translocation. (Scale bar 45 µm). (d, right) Bar graph showing the percentage of cells exhibiting β-arrestin-2 nuclear translocation in response to RTX (10 nM, 15 min) in the absence or presence of EGTA (10 mM) (RTX: 71 ±12.6, RTX+ EGTA 21.8 ± 13 P: * <0.0001). (e) Effect of PKC blockers on TRPV1-induced phosphorylation of ERK1/2 in HEK cells exposed to RTX for 5 min. (f) Densitometry analysis of pERK1/2 normalized to total ERK at 5 min post RTX and upon treatment with PKC blockers. *P <0.05. Data are expressed as means ± SEM. (g) Confocal image of TRPV1 HEK cells co-transfected with PKCβII-GFP and treated with RTX (10 nM). As indicated by the transient but pronounced translocation of the PKCβII-GFP from the cytosol to the plasma membrane, ligand-induced TRPV1 stimulation activates PKCβII. (h) Effect of β-arrestin2 knockdown on RTX-induced pERK1/2 probed by western blot.

### TRPV1 activation prevents MOR desensitization

To assess the functional consequence of β-arrestin2 nuclear translocation on MOR function and desensitization, we measured N-type (Cav2.2) voltage-gated calcium channel inhibition, as a surrogate of MOR-induced G protein coupling, following prolonged exposure to agonist. In control conditions, activation of MOR by DAMGO (1 µM) induces a rapid and robust inhibition of Cav2.2 N-type current in both transiently transfected HEK cells (47.23 ± 5.21 %, n=15) and native DRG neurons (55.57 ± 4.38 %, n=8) (**Fig. 6**). In contrast, absence of MOR or activation of TRPV1 alone does not promote calcium current inhibition (**Supplementary Fig. 7a**), confirming that agonist-bound receptors mediate the current inhibition. Following DAMGO pre-incubation for an hour, desensitized MOR does not modulate Cav2.2 currents, in both HEK (4.86 ± 2.83 % (n=9) and neurons (5.72 ± 2.42 % (n=7)) (**Fig. 6**), an effect relieved by a 60 min. wash out allowing recycling of internalized MOR (**Supplementary Fig. 7b**). We then tested whether activating TRPV1 rescues the N-type current inhibition by preventing β-arrestin-2 recruitment to MOR upon DAMGO exposure. As shown in **Fig. 6**, pre-incubation of cells with a combination of DAMGO and capsaicin (100nM) restores the MOR-driven N-type inhibition (51.29 ± 9.42 % (HEK cells n=9, **Fig 6a and b**) and 53.27 ± 4.5 % (neuron n=8**, Fig. 6c and d**), respectively), an effect lost in the absence of TRPV1 in both HEK cells **(Supplementary Fig. 7c)** and DRG neurons **(Fig 6d)**. Finally, MOR activation results in robust Cav2.2 inhibition in β-arrestin2^−/−^ HEK cells, despite pretreatment with DAMGO or DAMGO+capsaicin (47.93% ± 5.98, n=10; 47.11% ± 4.71, n=9; and 47.49% ± 5.04, n=10), supporting our hypothesis that β-arrestin2 translocation is essential in driving the TRPV1-induced blockade of MOR desensitization. Re-introducing β-arrestin2 to β-arrestin2-/- HEK cells re-establishes receptor desensitization and its relief triggered by TRPV1 activation (51.63% ± 4.66, n=9; 3.53% ± 1.56, n=10; 50.21% ± 2.17, n=10; respectively) **(Fig 6a and b)**.

**Figure 6.**
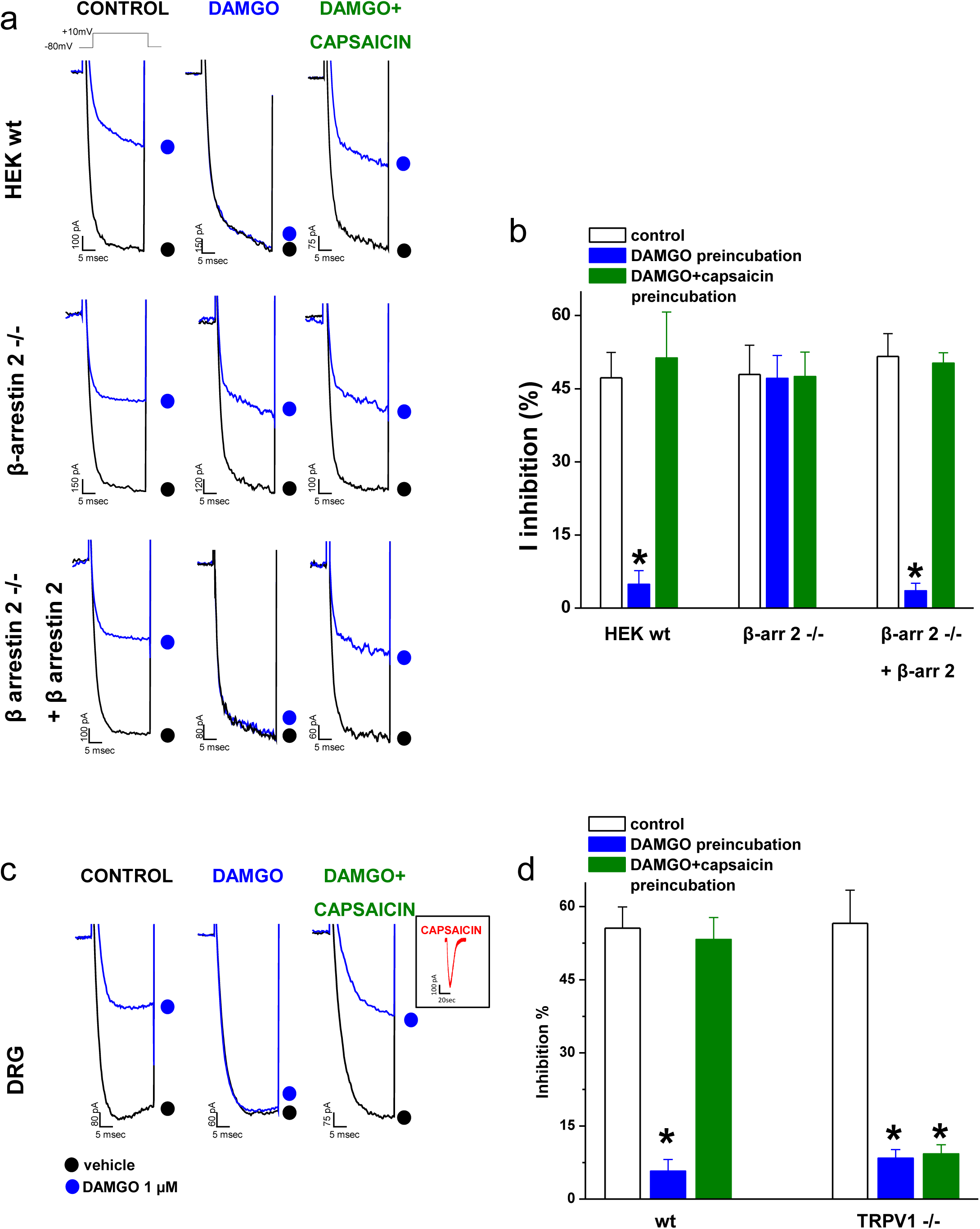
Activation of TRPV1 prevents acute desensitization of DAMGO-mediated inhibition of Cav2.2 current. (a) Representative calcium current traces (evoked at +10 mV from a holding potential of −80 mV during 30 ms) in HEK293 cells expressing CaV2.2, MOR1 and TRPV1, in the absence (black symbol) and presence (blue symbol) of 1 µM DAMGO. Middle pannel similar experiment as in upper panel using β-arrestin 2 -/- HEK cells. Lower panel shows that transfection of β-arrestin2 in β-arrestin2 -/- HEK cells is able to restore the inhibition of MOR desensitization mediated by TRPV1. The current was recorded in control conditions (left column), after 2 hrs pre-incubation with 1 µM DAMGO (middle column, acute desensitization) and after 2 hrs pre-incubation with 1 µM DAMGO and 100 nM capsaicin (right column). (b) Mean values of the peak current inhibition produced by DAMGO in untreated (white), DAMGO pretreated (blue) or DAMGO+capsaicin (green) pretreated cells for 2 hrs at 37° C (*n* values are, from left to right, of: 15, 9, 9, 10, 9, 10, 9, 10 and 10). (c) Representative native N-type current traces (recorded in the presence of Nifedipine 10 µM) in the absence (black symbol) and presence (blue symbol) of 1 µM DAMGO. Inset shows capsaicin-induced native TRPV1 current elicited at a holding potential of −80 mV by 100 nM capsaicin. (d) Mean values for the native Cav2.2 peak current inhibition produced by DAMGO application in the absence (white) and presence of DAMGO preincubation (blue) or DAMGO and capsaicin preincubation (green) in DRG neurons in wild type and TRPV1 ^−/−^ animals (*n* values are, from left to right, of: 8, 7, 8, 8, 6, 15). Data are expressed as mean values ± SEM. *P<0.01 (one-way ANOVA followed by Bonferroni post hoc test).

**Figure 7.**
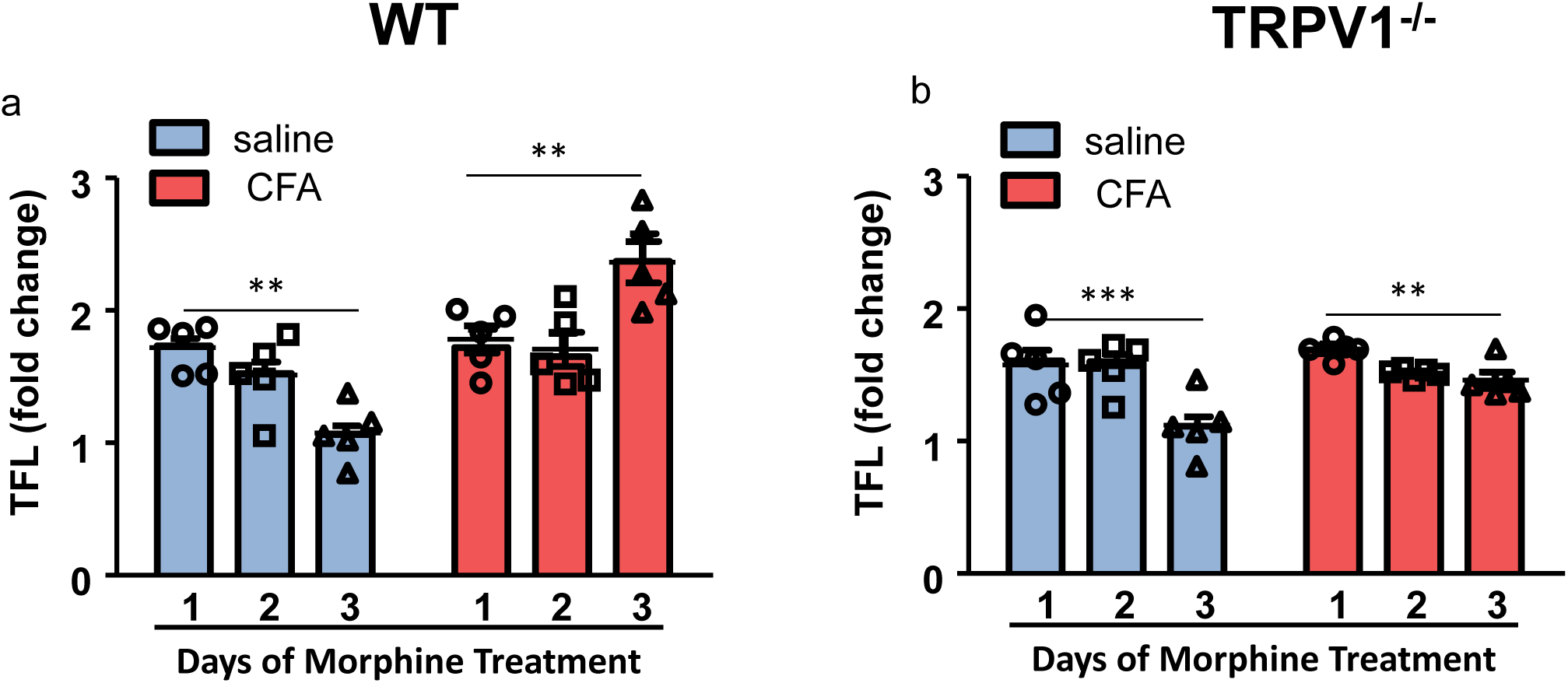
Activation of TRPV1 maintains exogenous opioid analgesia during inflammation. WT (a) or TRPV1^−/−^ (b) mice were treated with saline (blue) or CFA (red) and received Morphine-Sulphate (MS, 10mg/kg) twice daily for three consecutive days. Tail flick latency (TFL) before and 30 minutes after the treatment with MS was assessed daily, and the fold increase in TFL was calculated as a reflection of Morphine induced analgesia (WT-Saline-MS, n=5; WT-CFA-MS, n=5 / TRPV1-Saline-MS, n=5; TRPV1-CFA-MS, n=5). Results are expressed as mean±SEM and each symbol represent one mouse. Statistical analysis were performed using repeated two-way ANOVA followed by Tukey’s post hoc test (** p<0.01)

To determine whether TRPV1 contributes to the increased efficacy of exogenous opioids in inflammatory conditions^37^, we used an *in vivo* protocol of acute opioid receptor desensitization in which mice were treated with 10mg/kg of morphine i.p. twice daily for three days. This protocol causes a decrease in morphine efficiency measured with the tail flick latency test over the three day course of treatment in WT non-inflamed mice (**Fig 7 a, blue bar graphs**), and TRPV1^−/−^ non inflamed mice (**Fig 7 b, blue bar graphs**). We then assessed analgesia in morphine-desensitized WT and TRPV1^−/−^ mice subjected to CFA injection. Under these conditions, morphine analgesia was maintained in WT mice even after 3 days of repetitive morphine treatment (**Fig 7 a, red bar graphs**). This effect was dependent on the presence of TRPV1, as TRPV1^−/−^ mice show a decreased morphine analgesia in inflammatory conditions (**Fig 7 b, red bar graphs**). These results suggested that TRPV1 is responsible for maintaining opioid receptor signaling efficacy in inflammatory conditions. To address whether this facilitatory process occurs centrally or at the periphery, we assessed the dose-response of morphine analgesia after intrathecal or local injection, in CFA-inflamed mice, treated or not with morphine for three days **(Fig. 8a)**. Specifically, we compared morphine antinociceptive response in mice that were never exposed to morphine with mice that received 3 days of morphine injections. To determine morphine potency (ED50), these mice were given escalating doses of morphine via intrathecal (central) or local (peripheral) injection into the inflamed paw (**Fig 8**). When administered intrathecally in morphine-naïve mice, morphine dose-dependently attenuated CFA-induced mechanical hypersensitivity: the ED50 was comparable in WT and TRPV1-/- mice, suggesting that TRPV1 did not alter morphine antinociceptive potency in mice without previous exposure to morphine (**Fig. 8b and c**). In other words, acute morphine antinociception is not impacted by the absence of TRPV1. By contrast, following repeated morphine treatment, there was a notable reduction in morphine anti-nociceptive potency (i.e. increased ED50) **(Fig. 8b and c)**, suggesting decreased capacity of MOR to respond to morphine. When administered locally in the inflamed paw of saline treated CFA-inflamed mice, morphine dose-dependently reduced mechanical hypersensitivity **(Fig. 8d)**, thus confirming previous work that peripheral opioid receptors mitigate inflammatory pain under basal conditions^6,38,39^. However, as opposed to what is observed centrally, peripheral morphine potency is maintained in chronic morphine treated WT mice **(Fig. 8d and e)**. This result indicates that local inflammation prevents peripheral MOR desensitization upon chronic morphine treatment. Strikingly, this effect is dependent on the presence of TRPV1, as TRPV1^−/−^ mice exhibit a decrease in peripheral morphine potency (> ED50) as observed for centrally administered morphine **(Fig 8d and e)**. Thus, our findings show that TRPV1 is responsible for maintaining peripheral opioid receptor signaling in nociceptors in the setting of inflammation.

**Figure 8.**
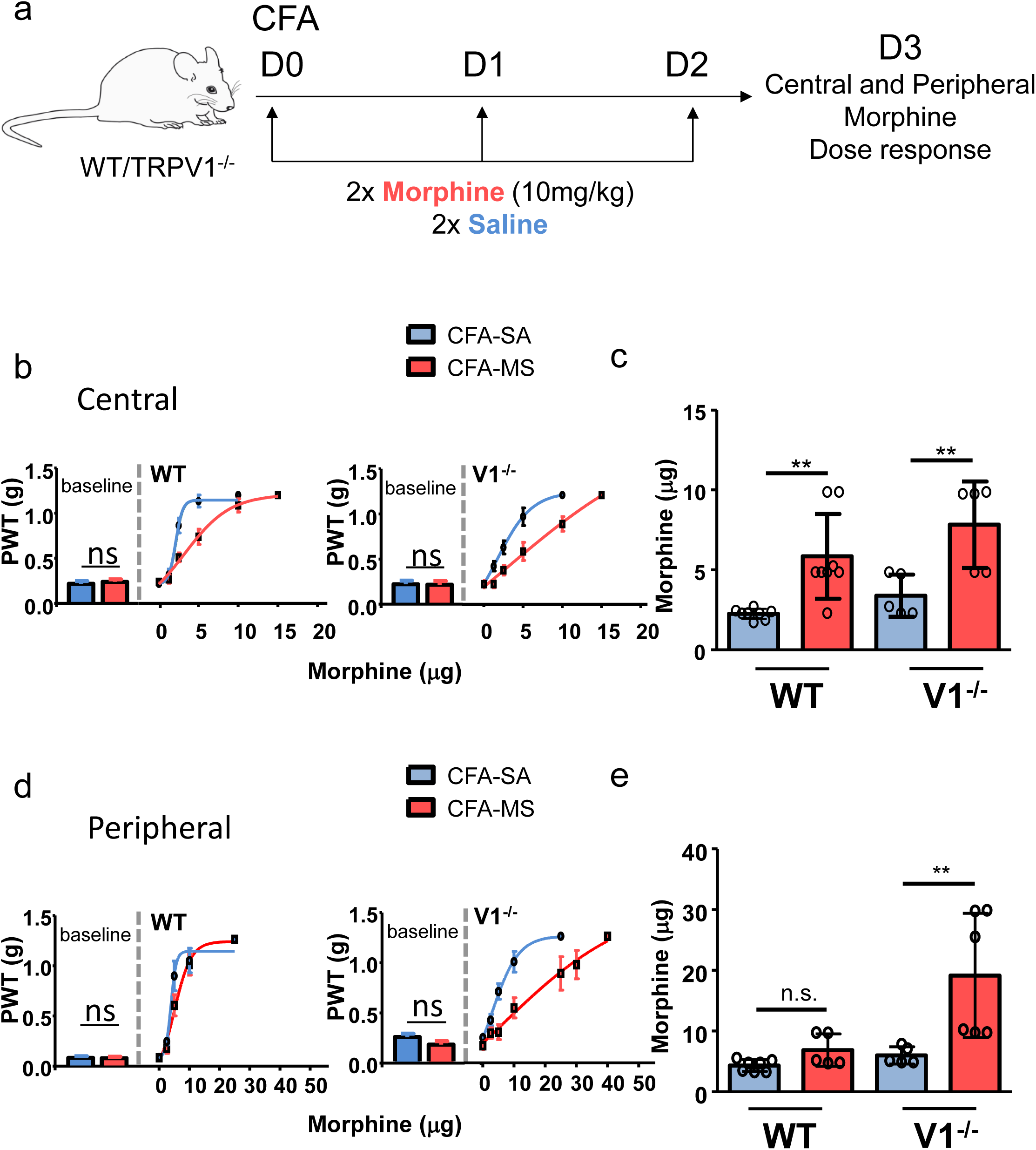
TRPV1 prevents peripheral opioid desensitization. (a) Schematic illustration of drug administration paradigm. Mice were treated with CFA (30μl; 1mg/ml) in the hind paw. On day 1, 2 and 3 of injection, they received either saline or MS (MS, 10mg/kg) twice daily. On day 4, potency of central opioid mediated analgesia was evaluated by measuring analgesia induced by intrathecal escalating dose of MS. Potency of peripheral opioid mediated analgesia was evaluated by measuring analgesia induced by local (ankle) escalating dose of MS. (b) Central MS dose„response curves in WT (left graph) and TRPV1^−/−^ (right graph) mice treated for 3 days with saline (blue line) or MS (red line). (c) Median effective dose (ED50) values as measured on day 4 after CFA treatment (WT-CFA-Saline, n=8; WT-CFA-MS, n=8 / TRPV1-CFA-Saline, n=5; TRPV1-CFA-MS, n=5). (d) Peripheral MS dose„response curves in WT (left graph) and TRPV1^−/−^ (right graph) mice treated for 3 days with saline (blue line) or MS (red line). (e) Median effective dose (ED50) values as measured on day 4 after CFA treatment (WT-CFA-Saline, n=7; WT-CFA-MS, n=5 / TRPV1-CFA-Saline, n=5; TRPV1-CFA-MS, n=6). Results are expressed as mean±SD and each symbol represent one mouse. Statistical analysis were performed using regular two-way ANOVA followed by Sidak’s post hoc test (** p<0.01)

Overall, our results suggest a model wherein activation of TRPV1 during an inflammatory insult translocates β-arrestin2 to the nucleus, which in turn prevents opioid-induced recruitment of β-arrestin2 to MOR and ensuing receptor desensitization, thus increasing opioid potency at peripheral nociceptors.

## Discussion

While opioids have been the gold standard of pain relief for decades, the current opioid crisis points to a lack of understanding of opioid receptor regulation in pathological conditions, particularly in the setting of peripheral inflammation. Earlier studies have proposed a positive modulation of opioid analgesia in inflammation^9,40^ whereas maladaptive processes occurring during resolution, combined with genetic and emotional predisposition, precipitate the transition to opioid insensitive chronic pain. Our findings shed light on the functional interplay between inflammation and opioid analgesia, along with the mechanisms that lead to acute and/or persistent receptor desensitization.

The main finding of our work is that activation of TRPV1 channels acutely results in the trafficking of β-arrestin2 from the cytosol to the nucleus. This channel-induced translocation attenuates the ability of β-arrestin2 to mediate MOR desensitization, thereby maintaining the sensitivity of MOR expressing nociceptors to the action of endogenously produced or exogenously administered opioids. Importantly, this mechanism of receptor regulation might be shared among other nociceptor-expressed GPCRs, like PAR2, localized in TRPV1-enriched membrane microdomains.

Of relevance to our new observations, a number of studies have highlighted TRPV1 as a key effector of MOR, supported by the fact that both are expressed in small unmyelinated DRG neurons^41^. Thus, on one hand, acute morphine administration can decrease capsaicin-induced TRPV1 current^42^ and reduce insertion of functional TRPV1 channels at the plasma membrane^43^ and, on the other hand, sustained morphine treatment may cause tolerance through the regulation of TRPV1 channel expression^44^. Here, our work reveals a reciprocal regulation of MOR by the TRPV1 channel and identifies the molecular underpinnings of this process involving the TRPV1-triggered translocation of β-arrestin2 to the nucleus. We found that TRPV1-/- mice are insensitive to endogenous opioid analgesia during resolution of inflammation. We then show that TRPV1 channel signaling overrides desensitization and maintains peripheral opioid receptor function, when using an *in vitro* or *in vivo* opioid receptor desensitization paradigm.

Agonists of the TRPV1 channel, including capsaicin containing cream have been extensively used as local analgesics. While Ca2^+^ dependent TRPV1 channel desensitization, inhibition of ion channels and nerve degeneration have been suggested to contribute to the analgesic effects of capsaicin, these mechanisms remain poorly defined, are reversible, and thus do not explain the long lasting pain relieving effects of capsaicin that last for weeks after treatment^45-47^. Previous work from Chen et al.^48^ determined that RTX-induced ablation of TRPV1-expressing afferent neurons prolonged opioid analgesia despite a reduction in MOR expression. Nevertheless, our findings indicate that even when keeping the integrity of the neurons, TRPV1 channel activity regulates β-arrestin2 mobilization to fine-tune opioid receptor function when it’s most needed. Additional work has described a direct inhibition of voltage-gated calcium channels by capsaicin in DRG neurons ^49-51^ which could contribute to decrease excitatory neurotransmission in the afferent pain pathway. We did not observe such effect in our hands when preincubating HEK cells or DRG neurons with DAMGO+capsaicin before wash out and recording N-type current inhibition. Furthermore, our BRET analysis suggests a shift in MOR signaling likely caused by an absence of β-arrestin2 interaction upon channel activation. Our electrophysiology data indeed indicate that Gβɣ modulation of N-type is restored by TRPV1 activation, yet a direct assessment of Gi/o coupling to MOR will ascertain that G protein biased signaling is favored upon TRPV1 channel stimulation. The main target of opioids used in the treatment of acute and chronic pain is represented by MOR and their effect is mediated partly by inhibition of high voltage gated N-type channels^52^. Sustained agonist activation of MOR, however, produces cellular opioid tolerance shown to be caused by β-arrestin2–dependent internalization of MOR which are trafficked to the endosomal compartments (for a review see^53^). It is well established that receptor desensitization and internalization are two processes that participate, among other mechanisms (resensitization, downregulation or de novo receptor synthesis) to opioid tolerance^54-56^. Therefore, further work will test this TRPV1-MOR paradigm in mouse models of opioid tolerance.

Finally our findings that β-arrestin2 translocates to the nucleus following channel activation will raise some new lines of research on the genetic and epigenetic mechanisms of pain modulation by this TRPV1-β-arrestin2 pathway. In contrast to previous studies showing physical association of β-arrestin2 with TRPV1, we could not find evidence of direct interaction between TRPV1 and β-arrestin2 using a BRET assay, or co-immunoprecipitation experiments (data not shown), and observation by confocal microscopy denotes distinct subcellular localization of the two proteins. Known as a central chaperone implicated in receptor signaling and trafficking, β-arrestin2 was already reported to shuttle from the nucleus to the cytoplasmic compartment, yet the presence of a NES in the 390–400 residues of the C-terminal domain excludes β-arrestin2 from the nucleus. Mutations of this motif or pharmacological inhibition of nuclear export were shown to promote translocation of β-arrestin2 into the nucleus^57,58^. Several potential roles of β-arrestin in the nucleus have been proposed, including transcriptional regulation, DNA repair or Histone acetylation^59,60^. The rapid and reversible trafficking of β-arrestin2 to the nucleus does not argue for apoptotic pathways to be engaged by the channel. However, we believe that transcriptional regulation mediated by TRPV1-β-arrestin2 signaling could contribute to the analgesic effect of topical capsaicin formulations used for pain management. Future proteomic and transcriptional studies may advance our knowledge of the factors interacting with β-arrestin2 as well as the genes regulated by TRPV1 signaling. It is likely that β-arrestin2 has an important function in the plasticity of nociceptive circuits post-inflammation.

Our new data emphasize information that appeared upon completion of our study, showing that TRPV1 activation blocks the opioid-dependent phosphorylation of the MOR by G protein-coupled receptor kinase 5 (GRK5)^61^. Thus, not only would TRPV1 activation attenuate the GRK-mediated recruitment of β-arrestin2 to trigger MOR internalization-desensitization, but the TRPV1-mediated translocation of β-arrestin2 out of the cytosol would also diminish its impact on MOR and likely other GPCRs present in nociceptors. Taken together, our work, along with the data of Scherer and coworkers, support a new concept that TRPV1 can bias MOR signaling via a dual mechanism involving the inhibition of receptor phosphorylation and the shuttling of β-arrestin2 to the nucleus.

Of note, we found that, although preventing calcium influx via TRPV1 blocks β-arrestin2 translocation, raising intracellular calcium by activating the calcium-permeant TRPA1 channel, or by using the calcium ionophore, ionomycin, did not stimulate the nuclear trafficking of β-arrestin2. Therefore, the dynamics of these events will have to be investigated further to understand the spatial and temporal requirements for the calcium signaling to trigger β-arrestin2 translocation, and to determine if there is co-trafficking of other signaling constituents (e.g. GRK5) along with β-arrestin2. To sum up, the rapid dual effect of TRPV1 activation to block GRK5 phosphorylation and to stimulate β-arrestin2 translocation can work together to leave the MOR in a sensitized state.

Overall, our work has unraveled a TRPV1-MOR interplay that governs opioid analgesia during inflammation. These results establish a novel framework for understanding the regulation of opioid receptor function in response to inflammation and the maladaptive processes that could lead to pathological pain during tissue healing. While clinical observations suggest that opioids are effective analgesics following acute injury, prolonged treatment for chronic pain conditions is often accompanied by side effects that may relate to MOR desensitization and the subsequent requirement of higher opioid doses to achieve the same analgesic effect. Our proposed model points to the TRPV1 as a molecular target, which could tailor strategies to enhance opioid efficacy early on while reducing tolerance. Therefore, if used in conjunction, TRPV1 agonists could improve the management of chronic and severe morphine resistant pain by high jacking endogenous mechanisms of receptor desensitization^62^, potentiating opioid analgesia and thus limiting dose escalation.

## Acknowledgements

We would like to thank Dr. Graciela Pineyro for technical assistance with the BRET assay and Michael Bruchas for his generous gift of the β-arrestin2-Venus and pRLuc8. This work was supported by operating grants from the Canadian Institutes of Health Research (CIHR) [CA, MDH, RT, TT], by the Vi Riddell Child Pain program of the Alberta Children’s Hospital Research Institute [CA, TT], and by the Fondation pour la Recherche Médicale (équipe FRM 2015) [EB] and the Agence Nationale pour la Recherche (ANR15-CE16-0012-01) [EB]. CA holds a Canada Research Chair in inflammatory Pain (Tier2). LB holds an ACHRI fellowship from the University of Calgary and a Cumming School of Medicine fellowship from the University of Calgary.

## Author contributions

CA conceived the study; LB, RA, MI, RF, EB, TT and CA designed the experiments; LB, RA, CYF, FA, RF and MI performed the experiments; LB, RA, CYF, FA, MI, RF and CA analyzed the data; LB, RA, MI, RF and CA wrote the paper with editorial input from MDH, EB, RT and TT.

## Material and Methods

### Mice

Six-week-old WT C57BL/6 mice were obtained from Jackson Laboratories (Bar Harbor, ME). TRPV1-/- mice (strain B6.129X1-Trpv1tm1Jul/J) were originally obtained from Jackson Laboratories (Bar Harbor, ME) and bred at the University of Calgary Animal Resource Center (RJ. Thompson). All mice were genotyped with the following primers: WT: cctgctcaacatgctcattg (984 bp); heterozygotes: tcctcatgcacttcaggaaa (450 and 984 bp); TRPV1-/-: tggatgtggaatgtgtgcgag (450 bp) (Jackson Laboratories). Transgenic Ai32/TRPV1-cre mice were bred in the University of Calgary Animal Resource Center (GW. Zamponi) by crossing mice homozygous for the Rosa-CAG-LSL-ChR2 (H134R)-EYFP-WPRE conditional allele (loxP-flanked STOP cassette) [strain B6;129S-Gt(ROSA)26Sortm32(CAG-COP4*H134R/EYFP)Hze/J; hereafter Ai32] with mice expressing cre recombinase in TRPV1 cells [strain B6.129-Trpv1tm1(cre)Bbm/J; hereafter TRPV1-cre] (both from Jackson Laboratories). All mice were housed under standard conditions with drinking water and food available ad libitum. All experiments were conducted on age-matched animals, under protocols approved by the University of Calgary Animal Care Committee and in accordance with the international guidelines for the ethical use of animals in research and guidelines of the Canadian Council on Animal Care.

### CFA model of inflammatory pain

6-week old C57BL/6 male mice were obtained from Jackson lab and were acclimated to the dynamic plantar aesthesiometer testing chamber for 2 days prior to starting the experiments. Mice were housed with free access to food and water, with a 12/12 light dark cycle. All experiments were conducted on age-matched animals, under protocols approved by the University of Calgary Animal Care Committee and in accordance with the international guidelines for the ethical use of animals in research and guidelines of the Canadian Council on Animal Care.

CFA was administered by intraplantar injection. All animals received 20 µL of CFA (1mg/ml, Sigma, St. Louis, MO), or 20 μL of saline as control, in the right paw. The animals were assessed daily on days 4-14 after CFA injections. Prior to nociception testing, mice were habituated to the behavior room for 1 hour. Mechanical threshold was assessed daily using the dynamic plantar test. Thermal sensitivity was measured using paw withdrawal latency to a radiant heat (Hargreaves apparatus from Ugo Basile). Briefly, mice on glass plates were subjected to targeted infrared heat under the ipsilateral or contralateral paw and the paw withdrawal latency was determined by motion-sensing software. Each time point was done in triplicate. Data are presented as mean values for each group ± SEM. Paw edema was measured throughout the experiment with a plethysmometer (Hugo Basile)

### *In vivo* opioid receptor desensitization paradigm

At day 0, mice were injected with CFA, or 20 μL of saline as control, in the morning. Morphine (10mg/kg, PCCA, London, ON) or saline (Sigma, St. Louis, MO) as control was injected intraperitoneally one hour after CFA, and then again in the afternoon. Twice daily morphine injections were repeated for two more consecutive days (day 1 and day 2). Tail flick latency to 46°C water bath was recorded before the CFA injection and on each day before and 30 minutes after the second injection of morphine. On day 3, mechanical nociceptive threshold were determined by the simplified up and down method using Von Frey filament in mice receiving increasing doses of Morphine injected either intrathecally (IT, 0;1;2,5;5;10;15 µg dose) or locally (in the ankle, 0;2,5;5;10;25;30;40 µg dose) in the inflamed paw.

### Plasmids

TRPV1-YFP and TRPV1-Rluc have been previously described^14^, TRPA1-YFP was constructed by cloning the TRPA1 coding sequence (gift from A. Patapoutian) into pEYFP-N1. TRPV1-and TRPA1-mCherry were produced by cloning the mCherry sequence in place of YFP in TRPV1- and TRPA1-pEYFP using (NotI and KpnI). TRPV1-CFP was generated by replacing the YFP coding sequence of TRPV1-YFP with CFP from pECFP (Clontech). β-arrestin2-YFP was a gift from Rithwik Ramachandran.

PAR2-YFP has been previously described^63^. Rat mu opioid receptor 1 (MOR1)-YFP was a gift from Gerald Zamponi and was used to generate MOR1-HA by cloning sticky-ended oligos containing the HA tag in place of YFP. MOR-Rluc8 was produced by using PCR to introduce AgeI and NotI sites to RLuc8 (a gift from Michael Bruchas) to clone in place of YFP. β-arrestin2-Venus was generated as previously described^64^. Protein kinase C-beta II-GFP (PKC-βII) was generously provided by Stephen S. G. Ferguson. β-arrestin-2 lentiCRISPR-V2 plasmids were a gift from Dr. Morley Hollenberg and contained guide sequences identified in a genome-wide screening. Humanized Renilla GFP^34^ was synthesized (Genscript) and cloned into pcDNA3.1. We added the SV40 large T antigen nuclear localization sequence PKKKRKVEDPKS^65^ by PCR, using an oligonucleotide containing the C-terminal 24 bp of rGFP, a 9 bp linker (GGTGGATCC) and the coding sequence of the NLS: CCGAAGAAAAAAAGGAAGGTTGAAGATCCGAAATCG. Nuclear localization of the resulting construct was confirmed by colocalization with DAPI. Constructs were confirmed with DNA sequencing.

### Cell culture and transfection

TsA-201 human embryonic kidney cells (HEK) were maintained in DMEM supplemented with 10% FBS, L-glutamine and penicillin/streptomycin at 37C in 5% CO2. Plasmids were transfected using calcium phosphate as previously described^66^ and β-arrestin siRNA (purchased from Santa Cruz) was introduced using Lipofectamine 2000. The p-ERK assay was done using 200 pmol of siRNA + 1 µg of TRPV1 transfected in a 35mm dish. Experiments were performed 24-48 hours after transfection.

For electrophysiology experiments using the β-arrestin2 CRISPR cell line, gene deletion was achieved by using published CRISPR guide RNAs (GAAGTCGAGCCCTAACTGCA and GCGGGACTTCGTAGATCACC,^67^) cloned into the lentiCRISPR V2 vector (Addgene #52961). CRISPR vectors were transfected into HEK cells using Lipofectamine LTX (Invitrogen) and cells were selected with 2.5 μg/ml puromycin. Successful β-arrestin2 knockdown was confirmed by western blot (**supplementary fig 4**).

### Western blot assay

Two days after transfection, cells were treated with agonist, harvested, pelleted and lysed in RIPA lysis buffer (1% Igepal, 0.1% SDS, 0.5% sodium deoxycholate in PBS) supplemented with a protease- and phosphatase-inhibitor cocktail (Complete Mini EDTA-free and PhosStop, respectively, Roche). Cleared lysates were separated by SDS-PAGE and transferred to nitrocellulose membranes. Membranes were blocked in 5% BSA (Sigma) or milk in TBS-T (50 mM Tris, 150 mM NaCl, 0.05% Tween-20. pH 7.5) and probed with the following antibodies: ERK, phospho-ERK, β-arrestin2 (C16D9), phospho-p38, p-38, phospho-JNK, JNK (all at 1:2000, Cell Signaling), GAPDH (1:1000, Santa Cruz Biotechnology). HRP-conjugated ECL-optimized secondary antibodies (GE Healthcare) were visualized on a Bio-Rad ChemiDoc. For p-ERK assays, cells were serum-starved for 3 hours.

### qPCR

Ipsi-lateral popliteal lymph nodes were harvested at day 3 and day 8 post CFA injection, dissociated using a bullet blender (Next Advance) with SSB05 beads (Next Advance) in RLT buffer (Qiagen, Toronto, Ontario, Canada). Total RNA was extracted using an RNeasy Mini kit (Qiagen), according to the manufacturer’s instructions. The quality and quantity of RNA were determined using a Nanodrop 2000c spectrophotometer (Thermo-Fisher Scientific, Montréal, Quebec, Canada). Relative Proenkephalin (PENK) gene expression (normalized to HPRT) was determined by qPCR using Quantitect SYBR Green PCR Master Mix (Qiagen) and a StepOnePlus real-time PCR detection system (Applied Biosystems, Burlington, Ontario, Canada). The following primers were used: 5′-GTTCTTTGCTGACCTGCTGGAT-3′ and 5′-CCCCGTTGACTGATCATTACAG-3′ for HPRT, 5′-CGACATCAATTTCCTGGCGT-3′ and 5′-AGATCCTTGCAGGTCTCCCA-3′ for PENK.

### FACS analysis

At day 8 of CFA, paw tissue was minced and digested in HBSS containing 1mg/ml of collagenase I (Roche) and 2.4U/ml Dispase (Gibco) for 90min at 37°C under agitation. After digestion, cells were filtered through a 70-mm mesh and washed in PBS 1% FBS. After blocking with CD16/CD32 (1/100 Fc Block, eBioscience) on ice for 15 minutes, cells were stained for 20 minutes with anti-mouse CD3-FITC (clone 145-2C11, BD Bioscience) and anti-mouse CD11b-PercP Cy5.5 (clone M1/70, eBiosicence), and analyzed on a FACS Canto II (BD Bioscience)

### Immuno-staining and confocal microscopy

Cells were plated on MatTek dishes coated overnight with poly-ornithine (0.002%), transfected with calcium phosphate and allowed to express for 48 hours before treatment. Fixed cells were permeablized with 0.1% Triton-X100 and probed with the β-arrestin2 antibody (Cell Signalling, 1:200); Images were obtained using a Fluoview (FV1000) laser scanning confocal microscope and a Zeiss LSM-510 Meta inverted confocal microscope.

### Chemicals and drugs

β-arrestin2 siRNA and control siRNA were purchased from Santa-Cruz Biotechnology. The TRPV1 agonists, RTX and capsaicin, were obtained from Sigma-Aldrich. The MAPK inhibitor (U-0126) was bought from Cell Signaling. The PKC inhibitor CGP53353 and GF109203X were purchased from Tocris bioscience. Ionomycin was obtained from Alomone Labs.

### Isolation of DRG neurons

DRG neurons were excised from 6 week old mice and enzymatically dissociated in Hank’s balanced salt solution (HBSS) containing 2 mg/ml collagenase and 4 mg/ml dispase (Invitrogen) for 45 min at 37 °C ^20^. DRGs were rinsed twice in HBSS and once in culture medium consisting of Dulbecco minimum essential medium supplemented with 10 % heat-inactivated fetal bovine serum (HI-FBS), 100 μg/ml streptomycin, 100 U/ml penicillin and 100 ng/ml Nerve Growth Factor (all from Invitrogen). Individual neurons were dispersed by trituration through a fire-polished glass Pasteur pipette in 4 ml media and cultured on glass coverslips, previously treated with HBSS with 25% Poly-Ornithin and Laminin (both from Sigma), overnight at 37 °C with 5% CO2 in 96% humidity. A subset of experiments were performed using DRG neurons obtained from TRPV1 knockout mice.

### MOR internalization assessment

HEK cells co-transfected with a DNA mix of TRPV1-mCherry, MOR-YFP, and were treated or not with DAMGO (1µM). In some conditions, cells were pre-treated with capsaicin (100nM) before DAMGO treatment. Cells were imaged on a Zeiss LSM-510 Meta inverted confocal microscope. Images were then analyzed with Image J software, using a sequence of events including background subtraction, transformation of MOR-YFP signal to an 8-bit image, adjusting threshold and noise reduction. Vesicles within the cells were then counted. The same sequence of events with the same settings was consistently applied to each analyzed image.

### BRET assay

BRET1 assay was performed between MOR-Rluc8 and βARR2-Venus, MOR-YFP and βARR2-Rluc^14^, or βARR2-RlucII and PAR2-YFP ^63^. Briefly, HEK cells were co-transfected with a DNA mix of (100ng donor:1µg acceptor) using lipofectamine. Cells were plated in a white 96-well plate (BRAND) for BRET assay. The agonist-induced BRET signal was calculated as the difference in BRET signal from cells treated with agonist or vehicle. For the dose-response BRET curve, cells were incubated with different concentrations of DAMGO for 15min at 37°C, then washed and treated with the RLuc substrate coelanterazine-h (5μM) for 5min prior to BRET measurement. BRET signal was measured in a Mithras LB940 (Berthold Technologies) and calculated as the ratio of YFP emission (530nm) to the RLuc emission at (460nm). BRET signal was expressed as net BRET, which is the difference between the signal from YFP/RLuc and the signal from RLuc alone.

For BRET assay between MOR-YFP and βARR2-Rluc, cells were transfected in 96 well plates using Lipofectamine 2000 (Invitrogen) with a DNA mix containing 100 ng of MOR-YFP plasmid (provided by Dr. S. Granier, IGF Montpellier) and 2 ng of Rluc-β-arrestin2 plasmid (kindly provided by Dr. M. G. Scott, Institut Cochin, Paris, France) and 40 ng of TRPV1 plasmid (obtained from Karel Talavera, Leuven Belgium) or empty vector. 24 h after transfection, BRET signal was measured using the pharmacological screening platform of the IGF, ARPEGE (http://www.arpege.cnrs.fr). Cells were washed twice with PBS and treated with coelanterazine-h (5μM) for 5 min. prior to stimulation with DAMGO (10 µM) or capsaicin (500nM), or DAMGO+capsaicin.

### Electrophysiology

Whole-cell patch-clamp experiments on HEK cells were performed 16-24 h after transfection with calcium phosphate^68^. The internal pipette solution contained (in mM):110 CsCl, 3 MgCl2, 10 EGTA, 10 HEPES, 5 MgATP, and 1 GTP (pH 7.2 adjusted with CsOH). The bath solution contained (in mM): 20 BaCl2, 1 MgCl2, 10 HEPES, 40 TEA-Cl, 65 CsCl and 10 D-glucose (pH 7.4 adjusted with TEA-OH) for HEK cells and 10 BaCl2, 10 HEPES, 120 TEA-Cl and 10 D-glucose (pH 7.4 adjusted with TEA-OH) for DRG. The bath solution for DRG recordings also contained 10 µM Nifedipine (Sigma, USA) to inhibit the native L-type calcium currents. Patch-clamp experiments were performed using an Axopatch 200B amplifier (Axon Instruments, Foster City, CA, USA) and pClamp 10.5 software was used for data acquisition and analysis (both from Molecular Devices Corp, Sunnyvale, CA). Data were digitized at 10 kHz and low-pass filtered at 1 kHz. Borosilicate glass (Harvard Apparatus Ltd., UK) pipettes were pulled and polished to 2-5 MΩ resistance with a DMZ-Universal Puller (Zeitz-Instruments GmbH., Martinsried, Germany). When applicable, voltages were corrected for liquid junction potentials. All experiments were conducted at room temperature (22 ± 2°C).

### Statistical analysis

Data analysis was completed using Clampfit 10.5 software (Axon Instruments). Graphs and statistical analyses were performed using Origin 7.0 analysis software (OriginLab, Northampton, MA, USA) for electrophysiology. Results were expressed as means ± S.E.M and numbers in parentheses reflect the number of cells (n). Statistical analyses were completed using either paired t-tests for all the results obtained before and after DAMGO in the same cells or unpaired t-tests when data were obtained in different cells. Where appropriate, analysis of variance (ANOVA) followed by Bonferroni’s post hoc test were performed. The criterion for statistical significance was set at p < 0.01 regardless of the method used.

## Supplementary Figure legends

**Supplementary Fig. 1. Peripheral endogenous control of inflammatory pain is also absent in female TRPV1 deficient mice.**

(a) CFA (30μl; 1mg/ml) was injected into the plantar surface of the right hind paw of WT (left graph) or TRPV1 -/- mice (right graph). Mechanical hypersensitivity was assessed by measuring paw withdrawal thresholds (PWT) using a dynamic plantar aesthesiometer. Mice were injected with either 10µl PBS (blue circle; n=6 for WT for and n=4 for TRPV1-/-) or Nal-M (2mg/ml, red square; n=6 for WT and n=4 for TRPV1-/-) in the ankle from day 4 to day 14, 30 minutes before mechanical pain assessment. Data are expressed as the mean ±SEM of PWT measured in the inflamed (CFA) ipsilateral paw (color) and contralateral saline-injected paw (grey) normalized to baseline. (* p < 0.05; two-way ANOVA followed by Bonferroni post hoc test). Area under the curve between day 6 and 10 were calculated and expressed as % of vehicle-treated animals for each genotype. Data are expressed as the mean min-max (* p < 0.001; Mann-Whitney U test) (b)

**Supplementary Fig. 2. Paw edema, T lymphocyte infiltration and opioid production is not altered in TRPV1-/- mice.** (a) Measure of paw edema in mice upon the different treatments indicated. (b) Measure of endogenous opioid expression in popliteal draining lymph nodes of CFA-treated WT or TRPV1^−/−^ mice at day3 and day8 of inflammation. Statistical analyses were performed using Mann-Whitney U test (*p<0.05) (c) FACS analysis of infiltration of paw by macrophages (CD11b) or T cell (CD3) at day 8 of CFA induced inflammation in WT and TRPV1^−/−^ mice.

**Supplementary Fig. 3. Activation of TRPV1 prevents MOR- and PAR2-β-arrestin2 interaction.**

Time course of agonist-induced BRET (DAMGO=10μM, capsaicin=500nM) between MOR-YFP and β-arrestin2-Rluc in absence a) or presence b) of TRPV1 channel. c) Quantification of the maximum BRET signal obtained in a and b. Data were normalized to DAMGO induced BRET and are represented as the mean± S.E.M Statistical analyses were performed using one way ANOVA and subsequent Dunett post hoc test (*p<0.001). Time course of agonist-induced BRET (2f-LIGRLO=10μM, capsaicin=500nM) between PAR2-YFP and β-arrestin2-Rluc in absence d) or presence e) of TRPV1 channel.

**Supplementary Fig 4. Generation of β-arrestin2 KO HEK cell line**

HEK cells were transfected with gRNA sequences cloned in the lentiCRISPR V2 vector, alone and together. β-arrestin2 deletion was assessed by western blot on whole cell lysate. GAPDH was used as loading control. Subsequent experiments were performed with the cells containing both gRNAs as the deletion was more complete.

**Supplementary Fig. 5. β-arrestin2 nuclear translocation is transient and induced by the specific activation of TRPV1 channel but not TRPA1**

(a) Nuclear localization of β-arrestin-2 following stimulation of TRPV1. HEK cells cotransfected with TRPV1 and β-arrestin-2-YFP were stimulated with 1µM capsaicin for 15 min, washed and fixed at the indicated time points. Percentage of cells in which β-arrestin2 was predominantly localized to the nucleus was quantified in each condition. Nuclear β-arrestin-2 in stimulated cells was significantly higher than that of unstimulated cells at the 15 and 30 min. timepoints (unstimulated, 2.7% ± 1.9; 15 mins, 48.8% ± 10.0, q=5.704; 30 mins, 33.4% ± 4.5, q=3.803, p<0.05, one-way ANOVA). b) Representative confocal image of HEK transfected with β-arrestin-2-YFP only and treated with RTX. Note the absence of nuclear translocation of β-arrestin-2. c) Representative confocal image of HEK co-transfected with β-arrestin-2-YFP and TRPA1-mCherry and treated with mustard oil (100μM) for 15 min.

**Supplementary Fig. 6. β-arrestin2 nuclear translocation after TRPV1 stimulation is associated with an impairment of MOR internalization upon DAMGO exposure.**

HEK cells transfected with β-arrestin2-YFP, TRPV1-CFP and MOR-mCherry were treated with RTX (10nM) and then with DAMGO (1µM) to activate MOR. Cells were then imaged using confocal microscopy.

**Supplementary Fig. 7. No respective effect of DAMGO or capsaicin are observed in absence of MOR and TRPV1.**

(a) Percentage of Cav2.2 current inhibition produced by DAMGO (black) or capsaicin (grey) in HEK cells transfected with CaV2.2 and TRPV1. (b) Percentage of Cav2.2 current inhibition produced by DAMGO on cells pre-incubated with DAMGO right after end of pre-incubation (solid bar) or after 30 or 60 minutes wash (open bars). (c) Percentage of Cav2.2 current inhibition produced by acute DAMGO on naïve HEK cells (white), after DAMGO pre-incubation (blue) or after DAMGO and capsaicin pre-incubation in HEK cells transfected with CaV2.2 and MOR (green).

